# glmGamPoi: Fitting Gamma-Poisson Generalized Linear Models on Single Cell Count Data

**DOI:** 10.1101/2020.08.13.249623

**Authors:** Constantin Ahlmann-Eltze, Wolfgang Huber

## Abstract

**Motivation:** The Gamma-Poisson distribution is a theoretically and empirically motivated model for the sampling variability of single cell RNA-sequencing counts (Grün *et al*., 2014; Townes *et al*., 2019; Svensson, 2020; Silverman *et al*., 2018; Hafemeister and Satija, 2019) and an essential building block for analysis approaches including differential expression analysis (Robinson *et al*., 2010; McCarthy *et al*., 2012; Anders and Huber, 2010; Love *et al*., 2014), principal component analysis (Townes *et al*., 2019) and factor analysis (Risso *et al*., 2018). Existing implementations for inferring its parameters from data often struggle with the size of single cell datasets, which typically comprise thousands or millions of cells; at the same time, they do not take full advantage of the fact that zero and other small numbers are frequent in the data. These limitations have hampered uptake of the model, leaving room for statistically inferior approaches such as logarithm(-like) transformation.

**Results:** We present a new `R` package for fitting the Gamma-Poisson distribution to data with the characteristics of modern single cell datasets more quickly and more accurately than existing methods. The software can work with data on disk without having to load them into RAM simultaneously.

**Availability:** The package `glmGamPoi` is available from Bioconductor (since release 3.11) for Windows, macOS, and Linux, and source code is available on GitHub under a GPL-3 license. The scripts to reproduce the results of this paper are available on GitHub as well.

---

The statistical distribution of sequencing counts from single-cell RNA-Seq is modelled by a Gamma-Poisson distribution

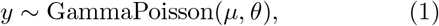

where *y* are the observed counts for a particular gene across a set of sufficiently similar cells (acting as replicates) and *μ* represents the underlying, true expression level of the gene: the expectation value, i.e., the value that would be obtained as the average of many repeated measurements. The parameter *θ* ≥ 0 determines the dispersion (width) of the distribution; the tightest case is *θ* = 0, in which case the distribution coincides with the Poisson distribution. Larger values of *θ* correspond to wider distributions.

Biological interest is added to (1) by extending it beyond (conceptual) replicates and letting *μ* itself vary across the cells. This can be done in different ways: via a generalized linear model, log *μ* = *Xβ*, as in the differential expression methods `edgeR` (Robinson *et al*., 2010; McCarthy *et al*., 2012) and `DESeq` (Anders and Huber, 2010; Love *et al*., 2014); via a factor analysis model (Risso *et al*., 2018); or via a matrix decomposition analogous to PCA (Townes *et al*., 2019). The fitted values of *β*, or the loadings of the principal components or the factors, then provide biological insight about “significant” variations in gene expression across cells, above and beyond the sampling noise.

A popular alternative approach is to transform the single-cell RNA-Seq sequencing counts using the shifted logarithm *f*(*x*) = log(*x* + *c*), with some choice of *c* > 0, and then proceed with analysis methods that are based on the least squares method, such as used for normal distributed data. However, this approach is fundamentally inferior as it overemphasizes the influence of small count fluctuations (Warton, 2018; Silverman *et al*., 2018) and deals poorly with variable sequencing coverage across cells (Townes *et al*., 2019).

With the Gamma-Poisson generalized linear model, parameter estimation proceeds by minimizing the deviance, a generalization of the sum of squares of residuals used in the least squares method. There are already a number of implementations to this end, including the R packages `MASS` (Venables and Ripley, 2002), `edgeR` and `DESeq2`. These all follow a similar approach: for each gene, the parameter vector *β* is estimated using an iterative reweighted least squares algorithm, and the dispersion *θ* is found by likelihood maximization. After years of development, and with tens of thousands of users, `edgeR` and `DESeq2` provide robust implementations and are a *de facto* standard for the analysis of bulk RNA-seq data.

Application of these implementations to single-cell RNA-seq data, however, suffers from several issues. First, their runtime becomes excessive as the number of cells gets large. Second, their functionality—fitting a GammaPoisson generalized linear model for a fixed, known design matrix *X*—is only part of what users need: with single-cell RNA-seq data, important research questions include identification of latent factors, dimension reduction, clustering and classification.

These limitations hamper the development and uptake of statistical models based on the Gamma-Poisson distribution and appear to be driving analysts towards the transformation approach.

The R package `glmGamPoi` provides inference of Gamma-Poisson generalized linear models with the following improvements over `edgeR` and `DESeq2`:

1. Substantially higher speed of the overdispersion estimation, by using an efficient data representation that makes uses of the fact that most entries in the count matrix are from a small set of integer numbers ({0,1, 2,…}).
2. More reliable estimation of the overdispersion on datasets with many small counts.
3. No size limitations for the datasets. `glmGamPoi` supports fitting the model without loading all data into RAM simultaneously (i.e. working “on-disk”), by using the `HDF5Array` (Pagès, 2020) and `beachmat` (Lun *et al*., 2018) packages.
4. Small number of dependencies to facilitate use as a building-block for higher-level methods, such as factor analysis, dimension reduction or clustering/classification.

The details of the algorithm are described in Appendix A.

Like `edgeR`, `glmGamPoi` also provides a quasi-likelihood ratio test with empirical Bayesian shrinkage to identify differentially expressed genes (Lund *et al*., 2012). In addition, it provides the option to form a *pseudobulk* sample, which Crowell et al. (2019) found to be an effective way to identify differential expression between samples for replicated single cell experiments.

To demonstrate how `glmGamPoi` can be integrated into other tools, we forked the `DESeq2` package and integrated `glmGamPoi` as an alternative inference engine (github.com/mikelove/DESeq2/pull/24).

We compared the runtime of `glmGamPoi` to other methods on the four single cell datasets summarized in Table 1. The timing results for the Mouse Gastrulation dataset are shown in Figure 1, those for the Brain20k, PBMC4k, and PBMC68k datasets are shown in Supplementary Figure S1. The speedup by `glmGamPoi` compared to `edgeR` and `DESeq2` was 6x to 13x. When the data were accessed directly from disk, the calculations took about twice as long. Omitting the estimation of the overdispersion parameter (by setting it to zero) reduced the runtime to about a half. The forked version of `DESeq2` that calls `glmGamPoi` was about as fast as calling `glmGamPoi` directly, indicating that inference carried out by `glmGamPoi` uses the largest part of the compute resources, while the additional steps carried out by `DESeq2` make relatively small demands. Although all methods theoretically scale linearly with the number of genes and cells, we find empirically that `glmGamPoi` scales better with the number of cells, which explains the observed performance benefit (Supplementary Figure S2).

**Figure 1:**
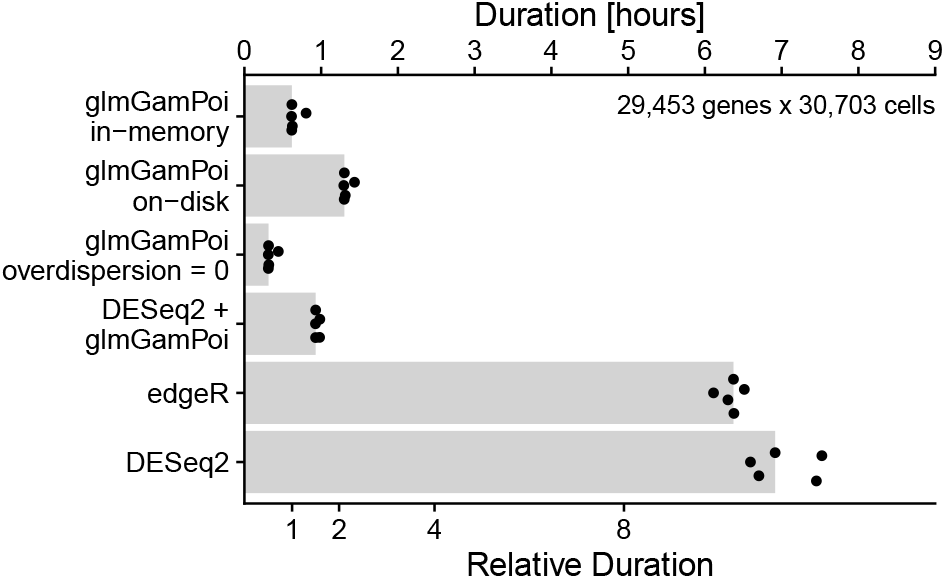
Bar plot comparing the runtime of `glmGamPoi` (in-memory, on-disk, and without overdispersion estimation), `edgeR`, and `DESeq2` (with its own implementation, or calling `glmGamPoi`) on the Mouse Gastrulation dataset. The time measurements were repeated five times each as a single process without parallelization on a different node of a multi-node computing cluster with minor amounts of competing tasks. The points show the individual measurements, the bars their median. Software versions: R 3.6.2, `glmGamPoi` 1.1.5, `edgeR` 3.28.1, `DESeq2` 1.29.4.

**Table 1:** Single cell datasets. In each, the number of genes was approximately 30,000.

| ID | Description | #cells |
| --- | --- | --- |
| Mouse Gastrulation | Effects of a gene-knockout on mouse gastrulation (Pijuan-Sala *et al.*, 2019). | 31,000 |
| Brain20k | Mouse brain atlas (10X Genomics, 2017a). | 20,000 |
| PBMC4k | Peripheral blood mononuclear cells (10X Genomics, 2017b). | 4,000 |
| PBMC68k | Peripheral blood mononuclear cells (10X Genomics, 2017b). | 68,000 |

On the PBMC68k dataset, the calculations of `DESeq2` and `edgeR` aborted because they ran out of memory (250 GB of RAM available). In contrast, `glmGamPoi` completed after ca. 45 minutes (Supplementary Figure S1).

Supplementary Figures S4–S6 show that `glmGamPoi`’s gain in performance does not come at a cost of accuracy. On the contrary, Supplement Figure S3 shows that `glmGamPoi` provides better estimates (in the sense of larger likelihood) than `DESeq2` for 72% of the genes and 10% of the genes in comparison with `edgeR`. Those differences with `edgeR`, seem to be of minor importance for assessing differential expression: the bottom rows of Supplementary Figures S5 and S6 show that the p-values from `glmGamPoi` and `edgeR` are very similar, consistent with the fact that they use the same statistical test. In Supplementary Figure S7 we provide a more detailed comparison for which genes the p-values of `glmGamPoi` and `DESeq2` are similar and for which genes they are different.

## Acknowledgment

We thank the authors of `edgeR` and `DESeq2` for developing and establishing the methods for handling high-throughput count data, on which this work relies. In particular, we thank Mike Love for his help in navigating the source code of `DESeq2` and explanations of underlying design decisions.

Furthermore we want to thank Will Townes, Mike Love, and Rahul Satija for their feedback on an earlier version of the software and the manuscript.

## A Methods

This section provides an overview over the implementation of `glm_gp`, the main function of the package `glmGamPoi`. `glm_gp` takes as input a count matrix and a specification of the design formula. In addition, it has several flags to turn on or off individual steps: size factor estimation, overdispersion estimation, overdispersion shrinkage, Cox-Reid adjustment. After validating its input, the function goes through the following steps and finally returns an estimate for the overdispersion for each gene, the regression coefficients for each gene and covariate, the deviance, the mean matrix and the size factors.

1. Estimate the size factors. `glmGamPoi` offers three different estimation techniques:

- Column mean
- Median of the ratios for each sample compared with a pseudo-reference sample (described by equation 5 in Anders and Huber (2010)). To avoid issues with the zeros, we calculate the ratio only on the positive counts.
- A deconvolution based method calling `scran’s calculateSumFactors` function The tuple of all size factor estimates is scaled by a common factor such that their geometric mean is one.
2. Estimate the overdispersion roughly using the empirical variances and the mean-variance relation of the GammaPoisson distribution *σ*^2^ = *μ* + *θμ*^2^.
3. Check if the experimental design is a single factor variable, in which case the coefficients of the GLM can be estimated for each group individually (Case 1). Otherwise, Case 2 applies.
4. Estimate coefficients for each gene roughly.

- Case 1: the estimates are the average counts per group.
- Case 2: fit the linear model.
5. Estimate the coefficients for each gene properly.

- Case 1: use a Newton-Raphson procedure to find the optimal coefficient for each group.
- Case 2: use Fisher-Scoring with a small (10^−6^) ridge penalty in combination with a line search to make sure each step decreases the deviance to find the coefficients for each gene.
6. Calculate the mean matrix *M* = exp (*BX^T^* + *D*), where *B* is the coefficient matrix, *X* is the design matrix, and *D* is the offset matrix.
7. Estimate the overdispersion for each gene properly by optimizing the log-likelihood of the Gamma-Poisson GLM. The optimization calls the `nlminb` function in `R` and uses the analytical expression of the first and second derivative to guide the optimization.

- If the first derivative is negative for *θ* = 10^−8^, we assume that there is no maximum and set the overdispersion estimate to zero
- We create a frequency table F = {{0, *f*_0_}, {1, *f*_1_}, {2, *f*_2_},…} for each count vector ***y***, where *f_k_* is the number of times that the count value *k* occures for that gene. We use this table to speed up parts of the log-likelihood calculation. For example, instead of calculating the log-gamma function for each element

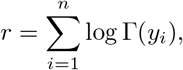

where *n* = length(***y***), we calculate

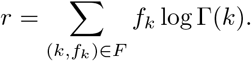 This optimization contributes most to the speed-up, because the table *F* is much smaller for many genes than the full vector ***y***. If we realize during the creation of the frequency table, that it grows too large and the additional cost of creating the table is not worth it, we revert back to the standard vector based optimization.
- The calculation of the first derivative is made more robust against numerical divergences by adding bounds on two terms of the sum.

– For small *θ*, the term 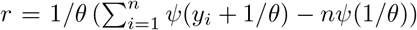 can become too large. We introduce an upper bound based on the Laurent series expansion of the equation at 1/*θ* = ∞ which is *r*_upper_ = Σ_*i*_ *y_i_* – (Σ_*i*_ (*y_i_* – 1)*y_i_*) *θ*/2.
– For small *μ_i_θ* the following term becomes imprecise 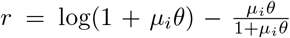. We use a Taylor expansion of log(1 + *μ_i_θ*) at *μ_i_θ* = 0 to derive an upper bound 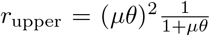 and lower bound 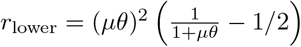 to stabilize the calculation.
8. Shrink the overdispersion estimates using the quasi-likelihood framework *σ*^2^ = *θ*_QL_(*μ* + *θ*_trend_*μ*^2^) that `edgeR` has developed (Lund *et al*., 2012).

- We fit a local median regression through the mean gene expression values and maximum likelihood overdispersion estimates *θ*_ML_ to identify the dispersion trend
- We convert the maximum likelihood overdispersion estimates to quasi-likelihood overdispersion estimates *θ*_QL_ using the relation 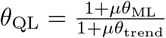.
- We estimate the parameters of the variance prior (the degrees of freedom df_0_ and scale 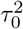 of an inverse Chi-squared distribution) using a maximum likelihood procedure.
- We shrink the quasi-likelihood overdispersion estimates by taking the weighted mean between the trended prior and the quasi-likelihood overdispersion estimates

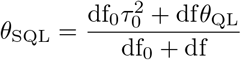

where *θ*_SQL_ is the shrunken quasi-likelihood overdispersion estimate.
9. Re-estimate the coefficients based on the updated overdispersion estimates.
10. Re-calculate the mean matrix based on the updated coefficients.

## B Supplementary Figures

**Suppl. Figure S1:**
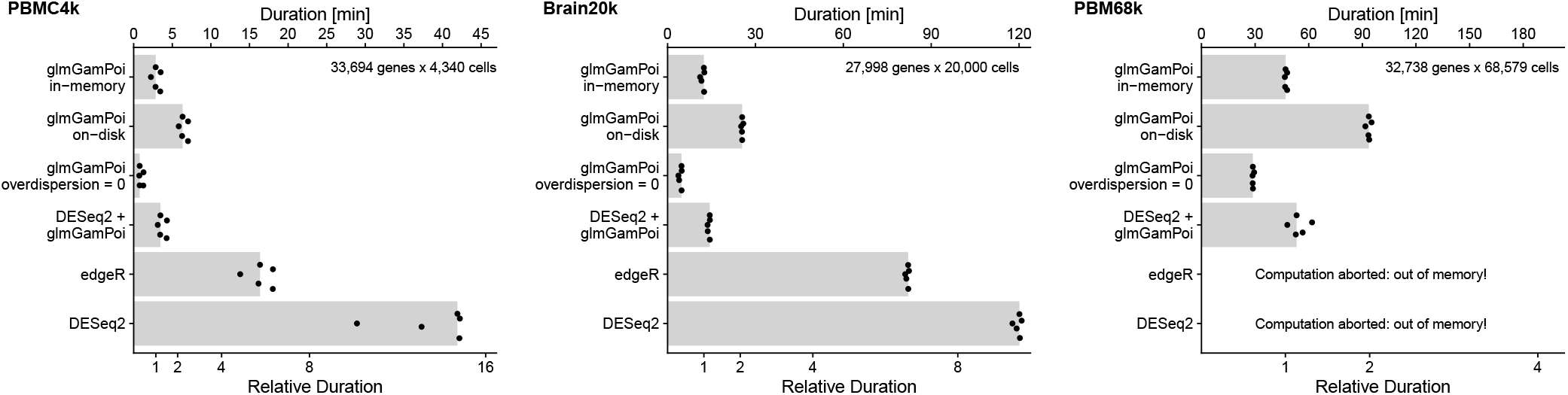
Extension of Figure 1 with three additional datasets: PBMC4k, Brain20k, and PBMC68k. edgeR and DESeq2 aborted the computation on the PBMC68k dataset because the available memory of 250GB was not enough. The runtime on the Mouse Gastrulation dataset is larger compared with the other three datasets, because a more complex experimental design was fitted. It took into account the mutation status of the *tomato* gene, the developmental stage, and the sequencing pool, which resulted in 5 covariates in the design matrix. The experimental design for the Brain20k dataset accounted for the mouse that was sequenced (2 covariates). For the PBMC4k and PBMC68k, we used an intercept-model (1 covariate).

**Suppl. Figure S2:**
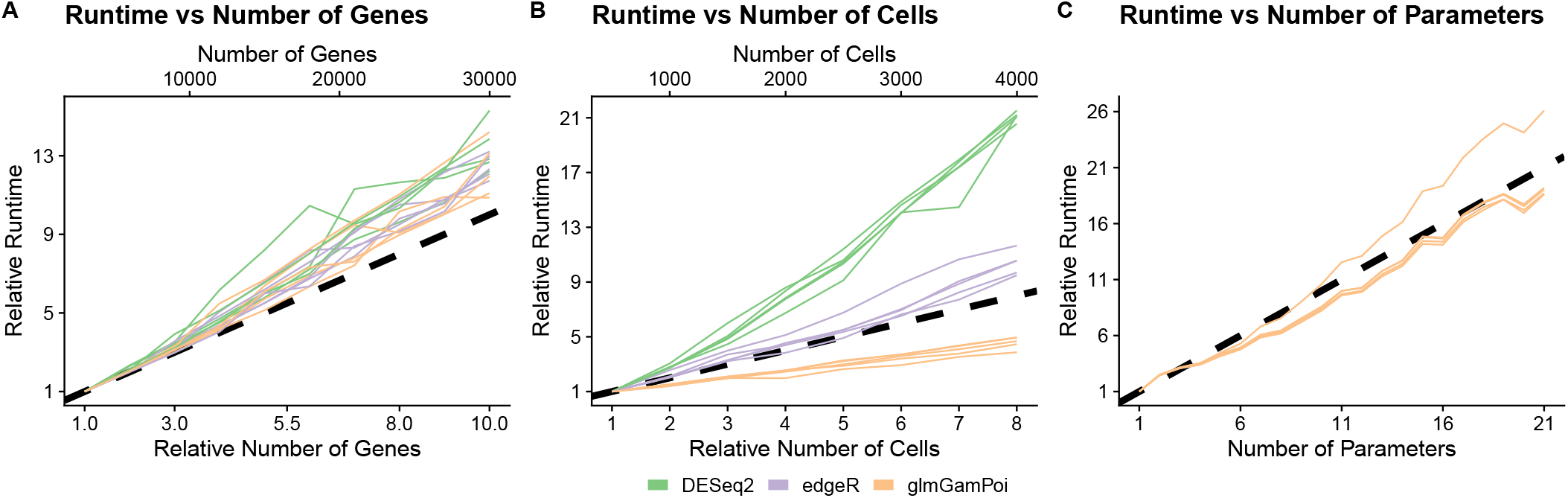
Line plots that show the increasing runtime of `edgeR, DESeq2`, and `glmGamPoi` with A) the number of genes, B) the number of cells, and C) the number of covariates in the experimental design. The dashed black line marks the diagonal which implies linear increase of the runtime. The experiments were run on different subsets of the PBMC4k dataset. The same methodology for measurements as in Figure 1 was used. Panel C) shows only the runtime for `glmGamPoi` because the runtime for `edgeR` and `DESeq2` is dominated by the overdispersion estimation, which masks the effect of the increasing number of parameters. Although asymptotically the scaling with the number of parameters is cubic in `glmGamPoi`, because of the QR decomposition of the design matrix, for the limited number of parameters that are typically encountered in real-world experiments, the scaling is well approximated by a linear function.

**Suppl. Figure S3:**
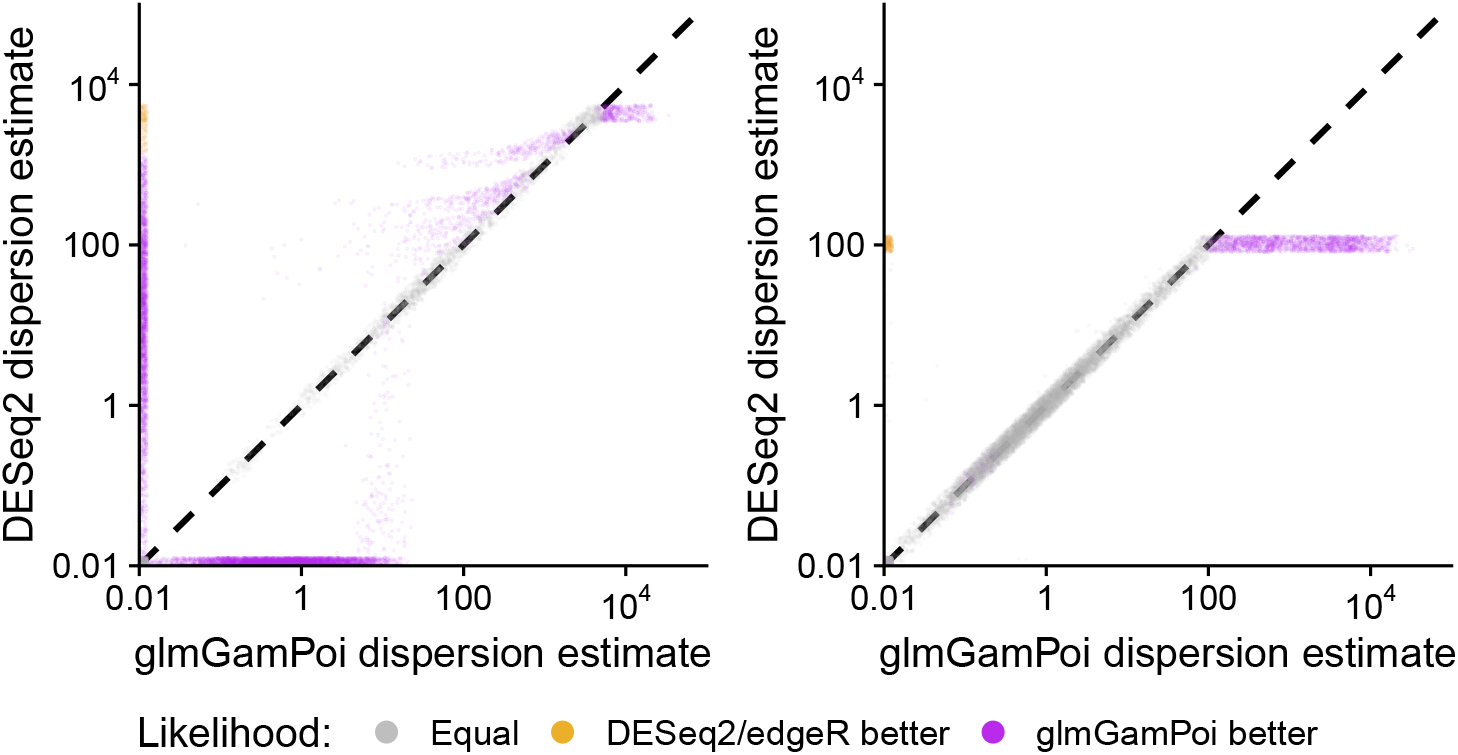
Scatter plots on a log-log scale that compare the genewise dispersion estimates of `DESeq2` and `edgeR` against `glmGamPoi`. A small jitter is added to each point to avoid overplotting. All methods optimize the same Cox-Reid adjusted Gamma-Poisson profile likelihood. This means that the best algorithm given the same data is the one that achieves the largest likelihood. The grey points show genes for which the likelihood was approximately equal (i.e. within ±0.001). glmGamPoi achieved a larger likelihood for 14,675 / 1,958 (DESeq2 / edgeR) genes (purple), an equivalent likelihood for 4,698 / 17,385 genes (grey), and had a smaller likelihood for 400 / 430 genes (orange). The dashed lines mark the diagonal of the plot.

**Suppl. Figure S4:**
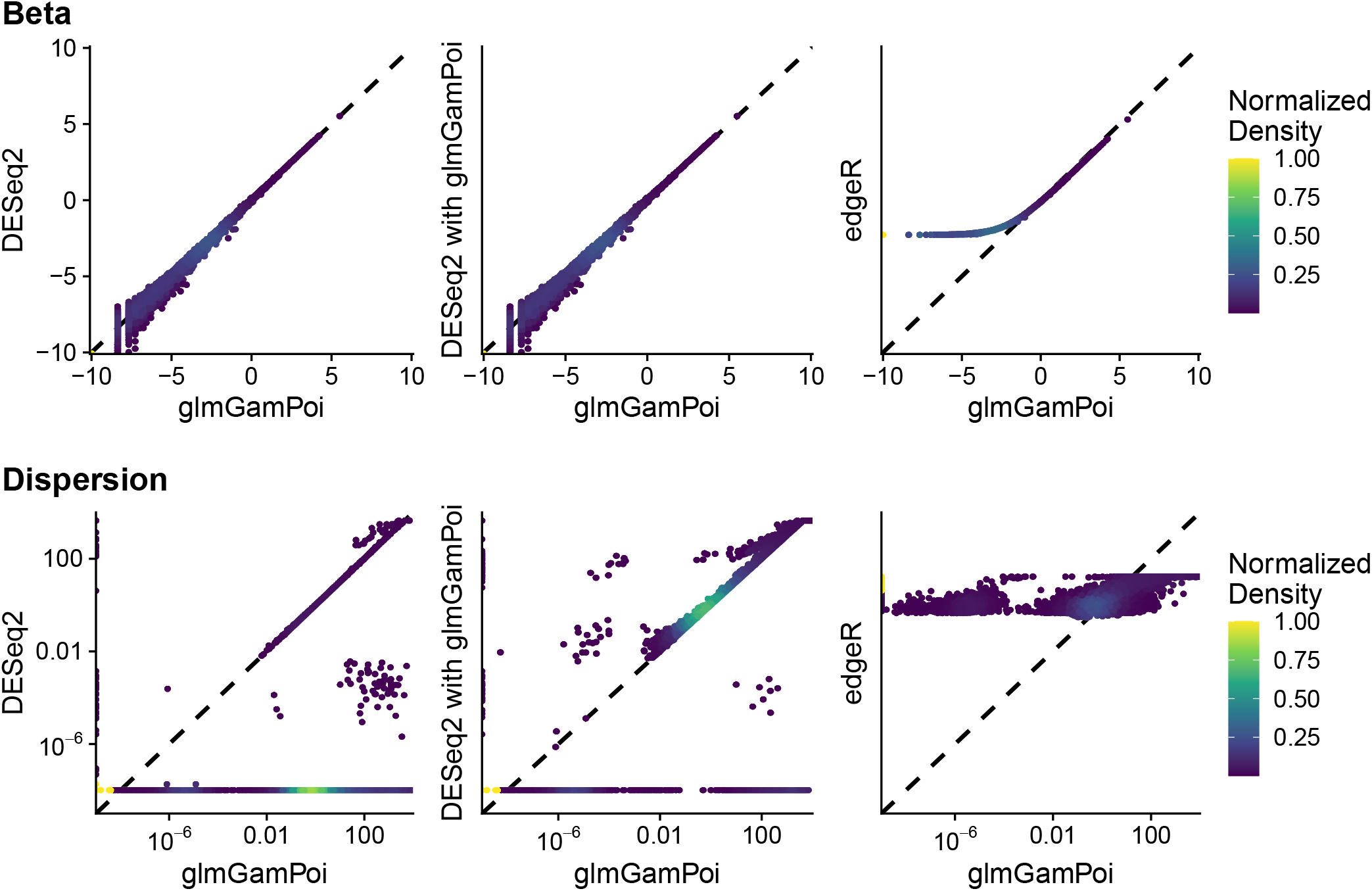
Scatter plots that compare the estimates of the intercept (one column in the Beta matrix) and the dispersion on the PBMC4k dataset for DESeq2, DESeq2 calling glmGamPoi, and edgeR against glmGamPoi. The color of the points shows the density of points in the area. The dashed diagonal line marks the diagonal where the two estimates are identical. The density for the color is normalized within each plot. Note that the dispersion estimates of `edgeR` are the shrunken gene-wise estimates. This is because `edgeR` does not provide a straightforward way to extract the maximum likelihood estimates.

**Suppl. Figure S5:**
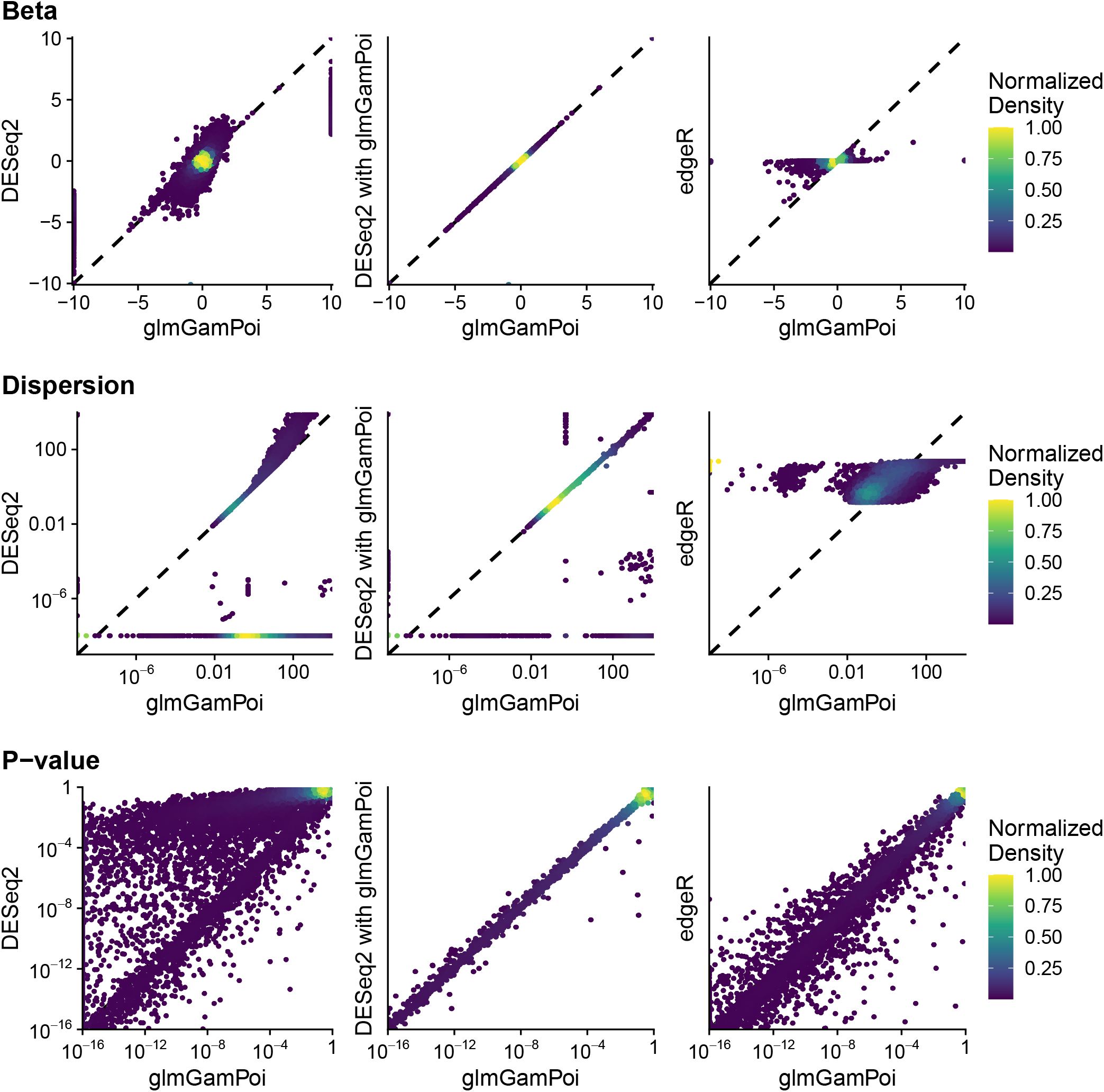
Scatter plots that compare the estimates of the *tomato* coefficient (one column in the Beta matrix), the dispersion, and the p-value from the test for differential expression on the Mouse Gastrulation dataset for DESeq2, DESeq2 calling glmGamPoi, and edgeR against glmGamPoi. Otherwise it has the same structure as Supplementary Figure S4.

**Suppl. Figure S6:**
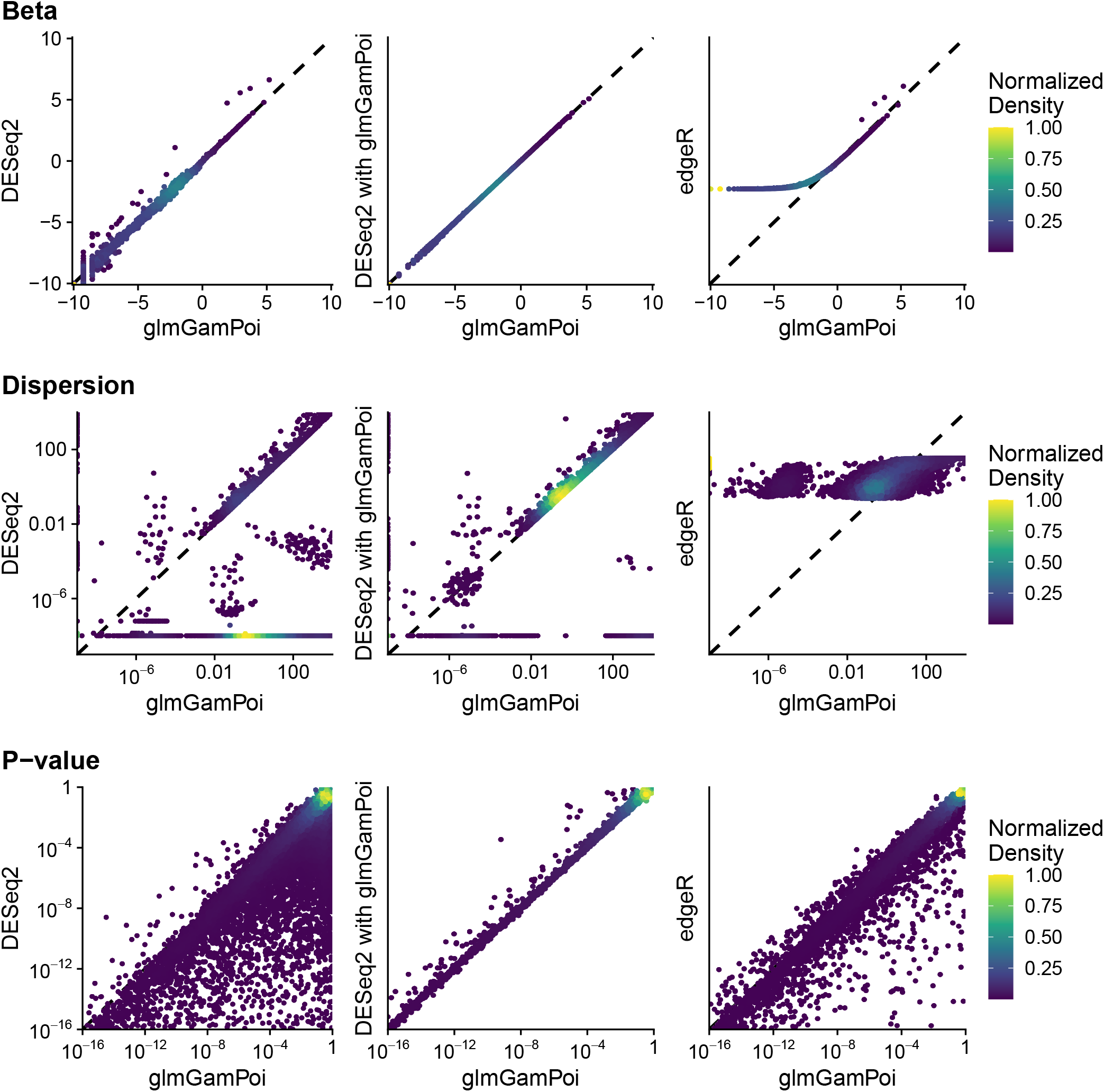
Scatter plots that compare the estimates of the *MouseA* coefficient (one column in the Beta matrix), the dispersion, and the p-value from the test for differential expression on the Brain20k dataset for DESeq2, DESeq2 calling glmGamPoi, and edgeR against glmGamPoi. Otherwise it has the same structure as Supplementary Figure S4.

**Suppl. Figure S7:**
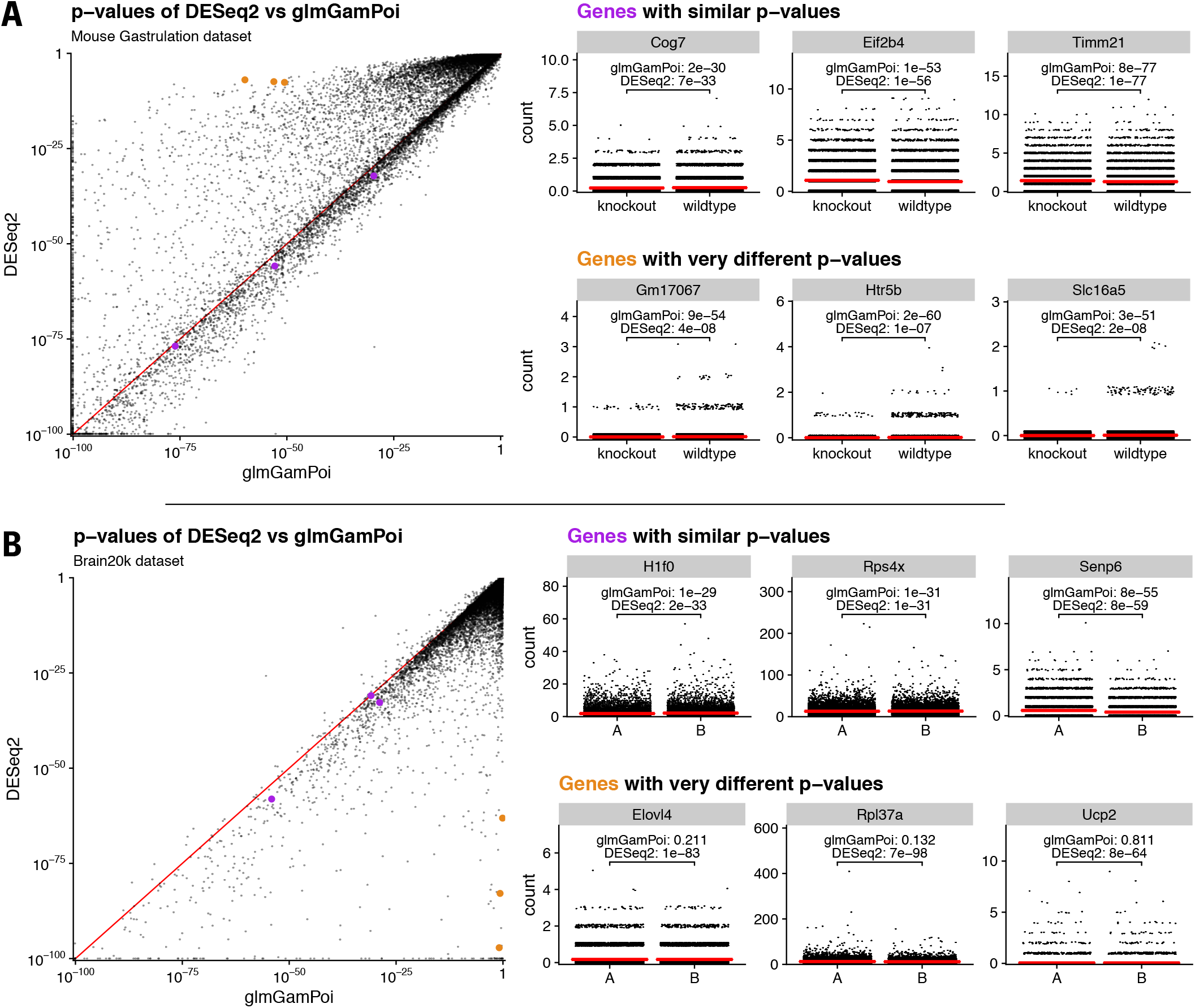
Comparison of the p-values calculated with the likelihood ratio test of DESeq2 and the quasi-likelihood ratio test of glmGamPoi. A) shows the differences on the mouse gastrulation knockout dataset. B) shows the difference on the Brain20k dataset. The scatter plots on the left show the p-values for the full vs. the reduced design on a double-logarithmic scale. Three exemplary genes with similar p-value are highlighted in purple and three genes with very different p-values in orange. Their counts are shown on the right. The red lines show per-group mean.

